# Path integration selectively predicts midlife risk of Alzheimer’s disease

**DOI:** 10.1101/2023.01.31.526473

**Authors:** Coco Newton, Marianna Pope, Catarina Rua, Richard Henson, Zilong Ji, Neil Burgess, Christopher T. Rodgers, Matthias Stangl, Maria-Eleni Dounavi, Andrea Castegnaro, Ivan Koychev, Paresh Malhotra, Thomas Wolbers, Karen Ritchie, Craig W. Ritchie, John O’Brien, Li Su, Dennis Chan, PREVENT Dementia Research Programme

**Affiliations:** Department of Psychiatry, University of Cambridge; Cambridge, UK; Cambridgeshire and Peterborough NHS Foundation Trust; Cambridge, UK; Wolfson Brain Imaging Centre, University of Cambridge; Cambridge, UK; Institute of Cognitive Neuroscience, UCL; London, UK; Jane and Terry Semel Institute for Neuroscience and Human Behavior, University of California; Los Angeles, USA; Department of Psychiatry, Oxford University; Oxford, UK; Department of Brain Sciences, Imperial College London; London, UK; German Centre for Neurodegenerative Diseases (DZNE); Magdeburg, Germany; Inserm, Institut de Neurosciences; Montpellier, France; Centre for Dementia Prevention, University of Edinburgh; Edinburgh, UK

## Abstract

The entorhinal cortex (EC) is the first cortical region to exhibit neurodegeneration in Alzheimer’s disease (AD), associated with EC grid cell dysfunction. Given the role of grid cells in path integration, we predicted that path integration impairment would represent the first behavioural change in adults at-risk of AD. Using immersive virtual reality, we found that midlife path integration impairments predicted both hereditary and physiological AD risk, with no corresponding impairment on tests of episodic memory or other spatial behaviours.

Impairments related to poorer angular estimation and were associated with hexadirectional grid-like fMRI signal in the posterior-medial EC. These results indicate that altered path integration may represent the transition point from at-risk state to disease onset in AD, prior to impairment in other cognitive domains.

## Main Text

Alzheimer’s disease (AD) is the leading cause of dementia and mortality, with limited treatment options (*1*). One major reason for the historic failure of therapeutic trials is the difficulty in identifying the first onset of clinically-relevant disease when interventions may have maximal value. While earlier detection of AD pathology is now possible with scalable tests for AD amyloid and tau biomarkers (*2*), these alone do not indicate the onset of clinical cognitive decline (*3*). Prevailing models of AD progression, which describe early damage to the entorhinal cortex (EC) and hippocampus within the medial temporal lobes (MTL), propose that cognitive changes are absent in preclinical AD (*4*). However, subtle deficits on bespoke MTL-supported cognitive tests are found in asymptomatic people at risk of AD who appear unimpaired on standard neuropsychological tests. Increased physiological risk of AD is associated with poorer allocentric spatial memory (*5*). Familial AD genetic risk gene carriers are impaired on a visual short-term binding paradigm (*6*), while carriers of the APOE-ε4 allele, the main genetic risk factor for sporadic AD, consistently perform worse on tests of path integration (*7–9*).

Path integration (PI) is a behaviour of high interest in early AD as it is thought to be subserved by spatially-modulated grid cells in the EC (*10*), the first neocortical region to exhibit tau pathology and neurodegeneration in AD (*11*). PI represents a form of navigation in which selfmotion cues are used to estimate environmental position. In an AD mouse model, EC tau pathology was associated with grid cell dysfunction and impaired spatial behaviour (*12*). Human studies have found that in patients with mild cognitive impairment (MCI) due to AD, PI error correlated with levels of cerebrospinal-fluid tau and EC volume (*13*), while in APOE-ε4 carriers, EC grid-like functional magnetic resonance imaging (fMRI) signal was reduced (*7*).

This study therefore tested the hypothesis that EC-related PI impairment represents the first cognitive manifestation of AD, with two predictions. First, that PI is impaired in people at risk of AD, independent of risk factor type. Second, that the PI deficit would occur prior to impairment in other cognitive domains potentially affected in preclinical AD.

Participants were recruited from the PREVENT-Dementia prospective cohort study (*14*) (n=100, 64 female, mean age 57 years, range 43-66, Table S1) and stratified according to three major late-onset AD risk factors: i) parental family history of dementia (n=62; FH), associated with a three-fold increased risk (*15*); ii) the Cardiovascular Risk Factors, Aging and Dementia Study (CAIDE) risk score derived from physiological variables including vascular health indicators, physical activity and education, associated with greater vascular pathology, tau accumulation and MTL atrophy over time (*16*); and iii) the APOE-ε4 allele (n=32), associated with 3-4 fold increased risk (*17*). Given that females have higher dementia prevalence (*18*), show diverging pathology progression in early AD stages (*19*), and that navigational strategies differ between males and females (*20*), participants were also stratified by sex.

PI was assessed using an immersive virtual reality (VR) paradigm with high sensitivity and specificity for prodromal AD (*13*). In an open-field environment viewed through a head-mounted VR, participants followed three cones in an outward L-shaped path before returning to their start location, thus completing a triangular trajectory (Fig. 1A). The return path was undertaken in three different environments to increase burden on self-motion cues: a) an unchanged “baseline” condition, b) a “no optic flow” condition, where surface information was removed to disrupt visual perceptions of movement speed, and c) a “no distal cues” condition where horizon orientation information was removed to disrupt visual perceptions of movement direction. The primary outcome measure was location error, i.e. Euclidean distance between participant estimate and true start cone location.

**Figure 1.**
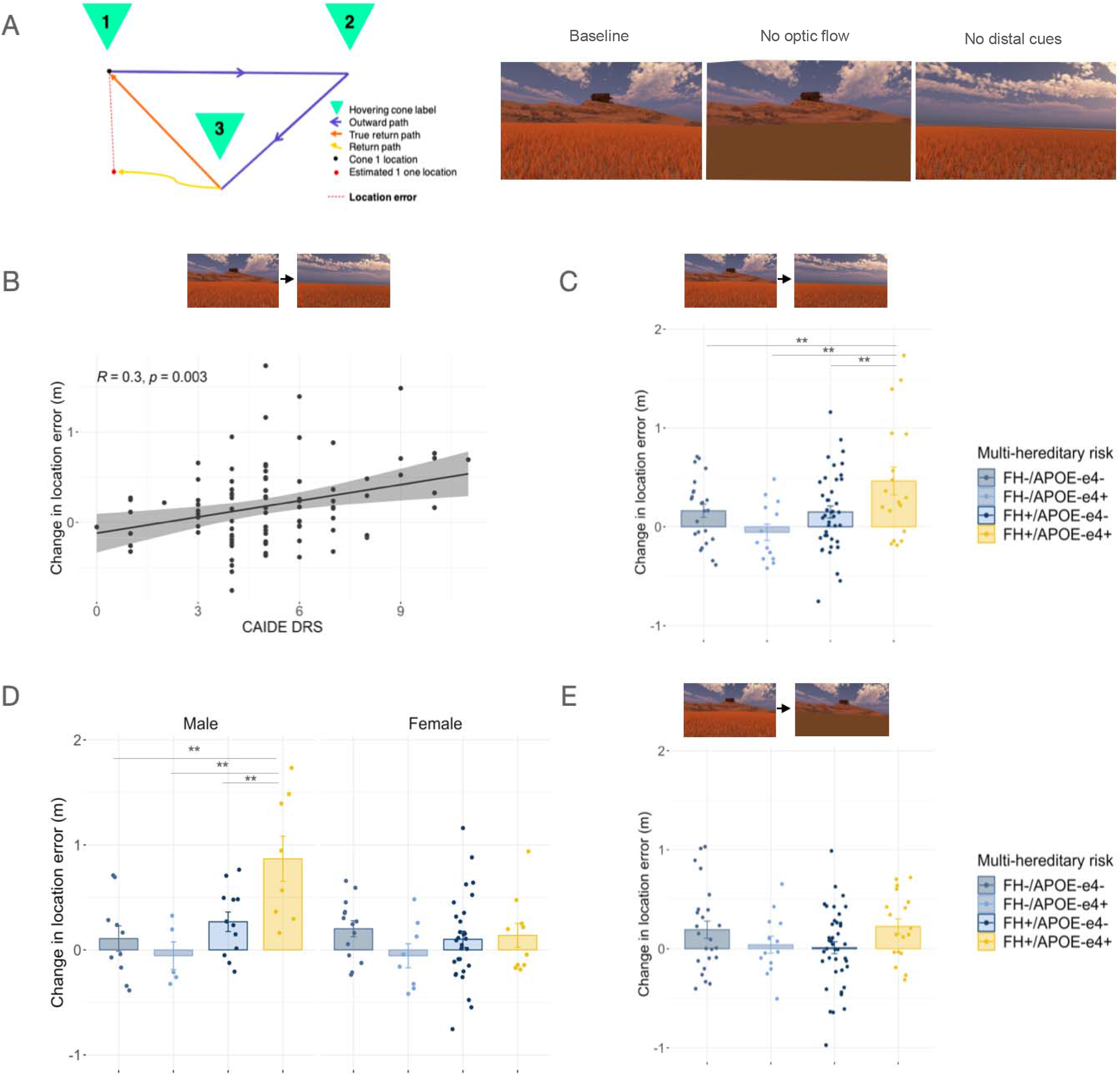
Hereditary and physiological risk factor PI impairments. (**A**): The PI task schematic and three return condition types. (**B**): Higher CAIDE dementia risk score significantly correlated with decline in performance on no distal cues condition relative to baseline in both sexes. (**C**): Decline in performance on no distal cues was largest in FH+/APOE-ε4+. (**D**): FH+ and APOE-ε4+ performance decline on no distal cues was specific to males (combined for display). (**E**): FH+/APOE-ε4+ or CAIDE (not pictured) decline on no optic flow condition relative to baseline was not significant. ** *p*_Tukey_ < 0.01

## Results

### PI impairments across risk factors

While we found no overall main effects of FH, APOE-ε4 or CAIDE score on location error when modelling individual trials with a mixed model approach (all *F* ≤ 1.17, *p* ≥ 0.283), several significant interactions between risk factors, return condition, and sex were present (all *F* ≥ 2.90, *p* ≤ 0.050). To study these further, we examined performance on the two manipulated return conditions relative to baseline (which additionally controls for individual differences in overall performance; Fig. 1A). Using linear regression to assess effects of all risk factors and sex together, we found significant worsening of PI performance after removal of the distal orientation cues across all individuals with elevated AD risk; namely, both a main effect of CAIDE (*F*_1,77_ = 11.04, *p* = 0.001; Pearson’s *r* = 0.30 *p* = 0.003; Fig. 1B) and two-way interaction effect of family history and APOE-ε4 status (*F*_1,77_ = 8.43, *p* = 0.005; hereditary x physiological interactions all *p* > 0.303). There were no effects of age or education (both *F* < 1.32, *p* > 0.254). Post-hoc analyses on this interaction showed that worsening performance was greatest in individuals with both FH+ and APOE-ε4+ (all t ≥ 3.28, *p_Tukey_* ≤ 0.008; Fig. 1C). Both FH and APOE-ε4 also interacted with sex (both *F* ≥ 8.11, *p* ≤ 0.006), with detrimental risk effects specifically occurring in males (Fig. 1D). We further explored the male-specific FH+ effect using estimated years to onset of dementia (EYOD; *18*), a temporal marker of preclinical state based on the difference between participant age and their parental age of dementia onset. In this cohort with a mean EYOD of 19.3 years, lower EYOD correlated with greater location error on the no distal cues condition (Pearson’s EYOD *r* = 0.55, *p* = 0.010; age *r* = 0.16, *p* = 0.300).

We confirmed these combined risk factor and sex-specific results by comparing the explained variance to simpler models (Table S2). The full model explained significantly more variance than the same model without a sex interaction (*F*_77,84_ = 3.05, *p* = 0.007; adjusted R^2^ = 0.26). It also explained more variance than separate models with individual risk factors (all *F*_77,90_ ≥ 2.84, *p* ≤ 0.002; all individual models adjusted R^2^ < 0.06), but not more than the same model with CAIDE omitted (F_77,85_ = 1.85, *p* = 0.081; adjusted R^2^ = 0.20), suggesting that the relationship between PI performance and AD risk was predominantly driven by hereditary risk factors.

### Decomposing PI impairments

In contrast to the “no distal cues” condition, performance differences between baseline and “no optic flow” conditions were borderline significant for hereditary but not physiological risk factors (two-way FH x APOE-ε4 *F*_1,77_ = 4.42, *p* = 0.039; post-hoc pairwise tests all *p_Tukey_* > 0.138; Fig. 1E), indicating that the PI impairment observed across all AD risk groups related specifically to orientation cue removal. To understand this impairment further, we decomposed location error into absolute distance and angular error (Fig. 2A; S1.4). Using the same risk factor x sex model, we found that orientation-related location errors in at risk individuals were driven by angular not distance errors, with the same main effect of CAIDE (angular error *F*_1,77_ = 10.42, *p* = 0.002; distance error *p* = 0.500) and two-way interactions of FH and APOE-ε4 together or individually with sex (angular error all *F*> 4.88, *p* < 0.030; distance error all *p* > 0.360). By wrapping the return angles of each trial to [−180,180] allocentric space, we determined that angular errors resulted from over-rather than under-turning (Fig. S1).

**Figure 2.**
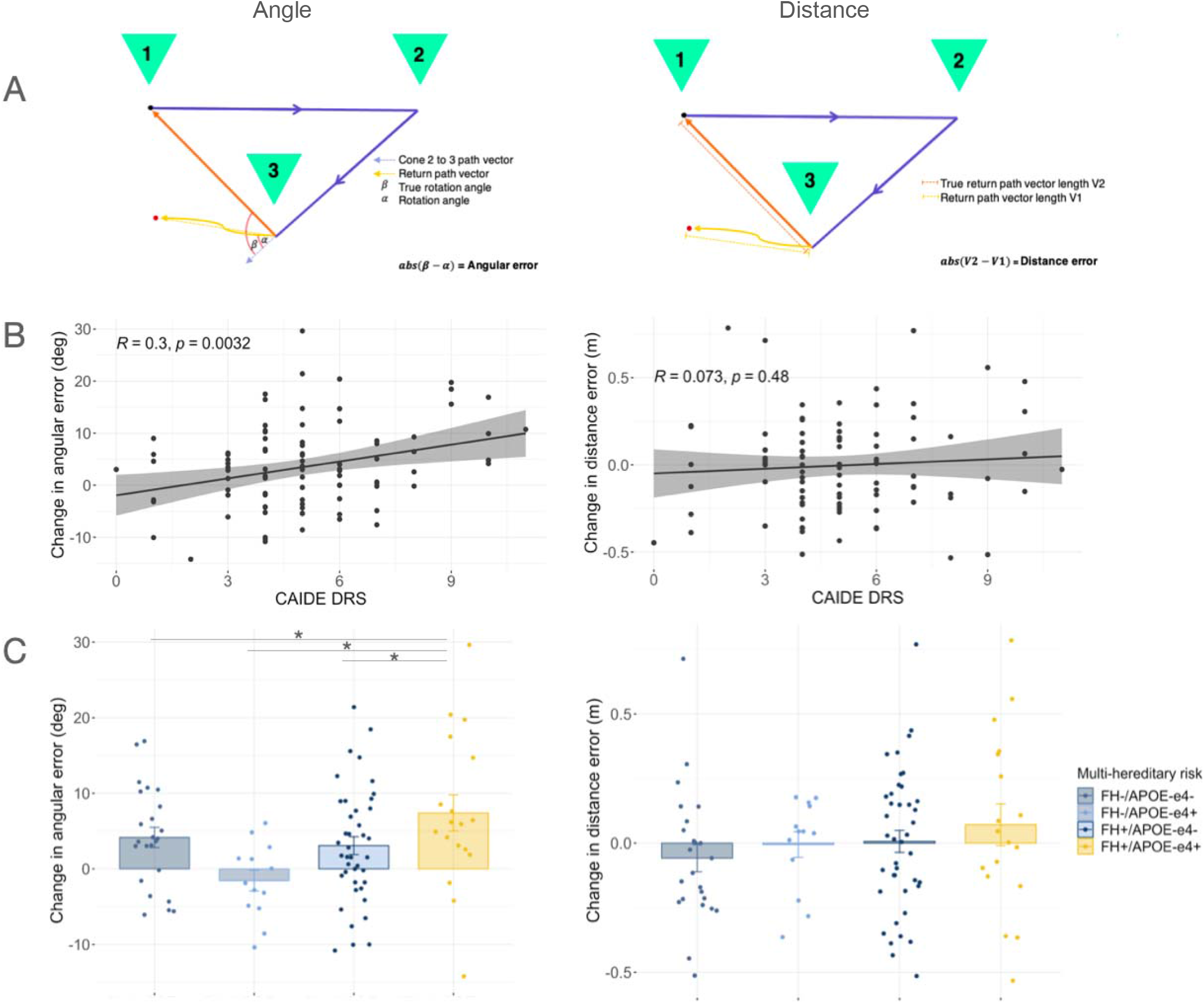
Angular rather than distance errors contributed to risk factor-associated impairment of PI after removal of distal orientation cues. (**A**): Schematic of angular error (left) and distance error (right) calculations. (**B**): CAIDE dementia risk score correlated with change in angular but not distance error from baseline to no distal cues. (**C**): Multi-hereditary risk interacted with sex for angular but not distance error changes in PI. \**p_Tukey_* < 0.05

### PI impairments are selective

For comparison, we tested other cognitive domains affected in preclinical AD (Table S3). Episodic memory, historically considered the domain first affected in AD, was assessed nonverbal name-face associative memory (*22*), visual short-term memory binding (*6*) and verbal narrative recall. Other aspects of spatial behaviour were tested with the Virtual Supermarket Trolley Task (VSTT) of egocentric spatial orientation (*23*), reflecting medial parietal lobe function and pertinent given early β-amyloid deposition in this region, and the hippocampusdependent 4 Mountains Test (4MT) of allocentric spatial memory (*24*).

We found no interactive effects of FH, APOE-ε4 or CAIDE on any comparator task (all *p* ≥ 0.14; Table S4). For episodic memory, no individuals at increased risk, regardless of risk factor, exhibited impairments on narrative recall or visual short-term binding (all *p* ≥ 0.100) while name-face association selectively correlated with CAIDE score (*F*_1,79_ = 21.28, *p* < 0.001; Pearson’s *r* = 0.41, *p* < 0.001). For spatial tests, only family history status had an effect, with FH+ individuals performing worse on the 4MT (*F*_1,74_ = 9.33, *p* = 0.003). When risk factors were separately modelled, female FH+ performed worse than FH-on the VSTT (*t*_92_ = 3.12, *p*_Tukey_ = 0.013; two-way interaction FH x sex *F*_1,92_ = 7.42, *p* = 0.008).

We compared the ability of the PI task to predict “double-risk” status (FH+/APOE-ε4+ vs any other combination) with that of the other cognitive tests with a cross-validated logistic regression using elastic-net regularisation to optimise the area under the curve (AUC) of the receiver operator characteristic. Though the mean AUC across 1000 iterations was only 0.67, 93% of iterations included a non-zero contribution from PI performance, whereas < 1% of iterations included a non-zero contribution from any other cognitive test. When age, sex, or education were used as predictors, the mean AUC was 0.56. In summary, PI was the only behaviour predictive of multifactor hereditary AD risk.

### Structural MRI correlates of PI impairments

In a subset of 55 participants (Table S5), ultra-high field 7T MRI was used to assess the volume of brain regions of interest (ROIs) associated with both navigation and early AD, namely the EC-hippocampal subfields, retrosplenial cortex and posterior cingulate gyrus (*25*). There were no significant differences in regional volumes between high and low risk participants across individual or interacting risk factors after multiple comparison correction (all *p*_FDR_>0.183; Table S6). When predicting change in performance from baseline to no distal cues, after adjusting for age, education and sex, there was no effect of any ROI independently, or interaction between ROI and risk factors (all *p*_FDR_ > 0.360), or even a combined effect when adding all ROIs into a single linear model (*F*_14,37_ = 1.39, *p* = 0.208).

### Functional MRI correlates of PI impairments

We employed a spatial memory paradigm found to elicit grid-like fMRI signals in the EC (*26*; Fig. 3A). We focused on the right posteromedial EC, the human homologue of rodent medial EC where most grid cells are resident and where fMRI signal was reduced in APOE-ε4 carriers (*7*) and healthy older vs younger adults (*26*). Adopting a similar analysis procedure with partitioning by odd/even grid events (*27*), we found that hexadirectional grid-like fMRI signals were not significant in the population overall (one-way t-test t_52_=0.50, *p*=0.600), but individuals with greater signals showed smaller PI performance declines from baseline to no distal cue conditions (β = −0.27 ± 0.10, *t*_46_ = 2.66, *p* = 0.011; Fig. 3B). Risk factors alone had no effect on the signal magnitude (all main and interaction effects *p* > 0.131). However, PI performance was predicted by a two-way interaction between CAIDE and grid-like signal (*F*_1,45_ = 4.20, *p* = 0.046; Johnson-Neyman interval for CAIDE > 7 β = −0.47 ± 0.15, *t*_45_ = 3.17, *p* < 0.001; no hereditary x CAIDE risk interaction; Fig. 3D) and by three-way interactions between individual hereditary risk factors, sex and grid-like signal (FH interaction *F*_1,41_ = 4.16, *p* = 0.048; APOE-ε4 interaction *F*_1,41_ = 3.52, *p* = 0.068; Fig. 3C). Including grid-like fMRI signal as a predictor in these models provided a better fit for the PI behavioural data than null models without fMRI inclusion (all *F* > 3.42, *p* < 0.017; for controls see S2.2), suggesting that impaired PI performance across risk groups was associated with altered grid-like fMRI signal in the posteromedial EC. More specifically, the higher risk individuals with poorer PI performance showed negative hexadirectional grid-like activity magnitudes. Negative magnitudes related to grid-like signal drift over time have been reported in healthy individuals and APOE-e4 carriers with poorer path integration ability (*7, 8, 26*). However, negative grid magnitudes due to temporal signal drift were not consistent with our data partitioning procedure, which used interleaved odd/even grid events to create estimation and test data sets. We hypothesised that non-6-fold symmetries might have contributed to the negative grid magnitudes after observing unbalanced directional sampling between partitioned sets, which we explored in a supplementary analysis (S2.3). While there were no effects of 4, 5, or 7-fold symmetries on PI performance (all *p* > 0.270), we found that a unidirectional signal consistent with head direction like processing predicted poorer PI performance after removal of distal orientation cues (β = 0.31 ± 0.09, *t*_46_ = 3.50, *p* = 0.001; Fig. 3E), complementing in reverse the sex and AD risk effects observed with the hexadirectional grid-like activity (Fig. 3C and 3F).

**Figure 3.**
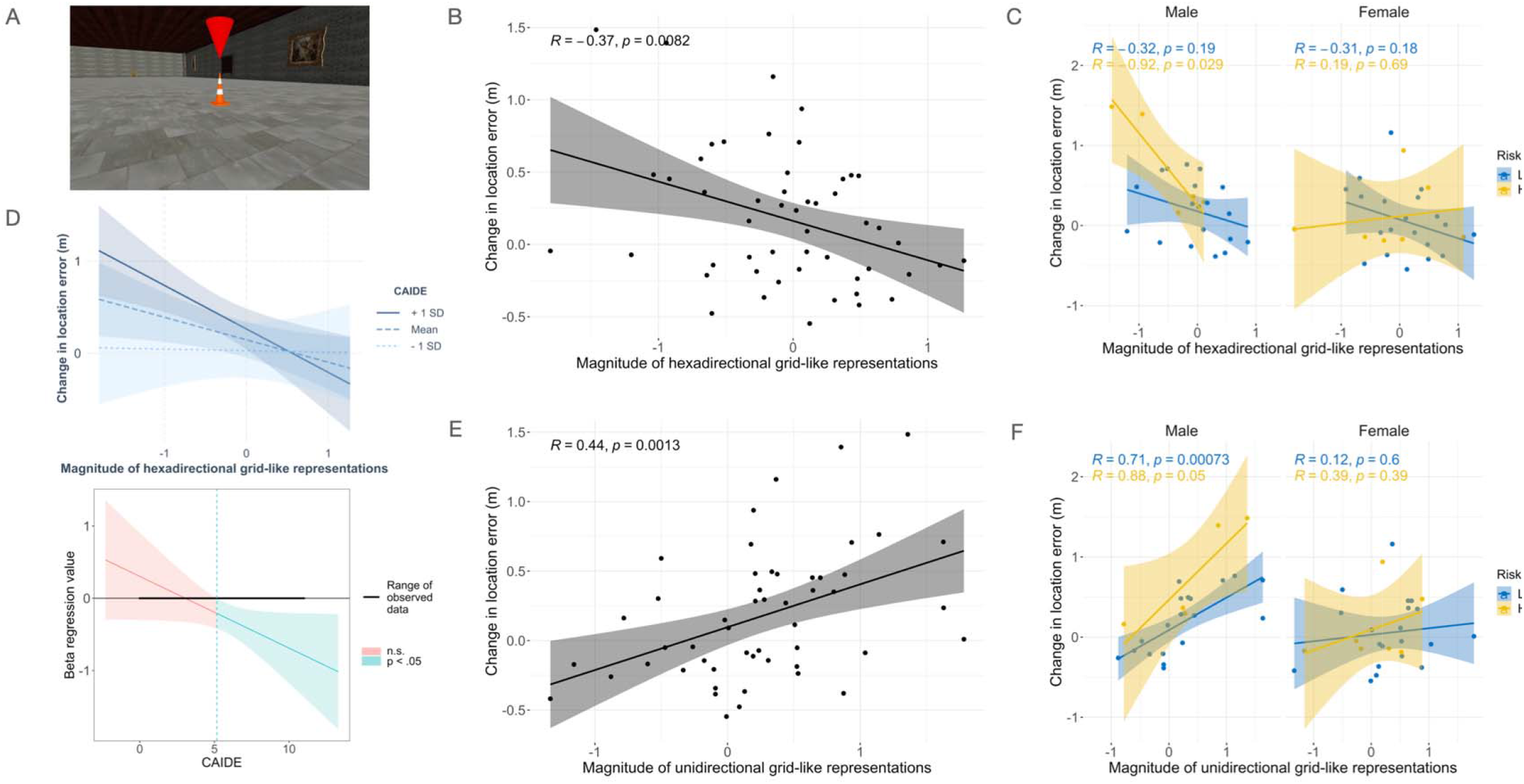
Right posterior-medial entorhinal fMRI correlates of decline in path integration performance. (**A**): Image of object memory location fMRI task. (**B**): Greater change in location error from baseline to “no distal cues” return condition associated with lower 6-fold grid-like activity, which reflects behavioural findings of showing a stronger effect in males with hereditary risk (**C**) or individuals with a higher CAIDE score (**D**). High risk is defined as FH+/APOE-ε4+ and low risk as individual or no FH/APOE-ε4 risk factors. (**E**): The negative 6-fold grid magnitudes in (**B**,**C**) might relate to a unidirectional head direction-like signal, which was stronger in individuals that showed worse PI in the absence of orientation cues. Again, this appeared specific to males with hereditary risk (**F**) in an exploratory analysis.

## Discussion

These results show that impaired PI selectively predicted both hereditary and physiological AD risk in asymptomatic middle-aged adults. We found no similar effect for other assessments previously reported to either show preclinical AD cognitive changes or probe the function of other AD ROIs. These included a test of visual short-term memory binding from presymptomatic familial AD cohorts, and tests of episodic memory, the mainstay of current AD cognitive assessments worldwide.

Our findings mirror neuropathological data implicating early AD pathology in the EC, which disrupts neuronal activity and spatial memory in animal models (*12*). The specific deficit after removal of orientation cues when only self-motion cues are available is consistent with grid cell stability being dependent on environmental boundaries (*28*) - imitating PI deficits in APOE-ε4 carriers (*7, 8*). Our additional observation that defective angular estimation drove PI errors is also in line with increasing evidence that spatial navigation is underpinned by neuronal vector-based coding (*29, 30*). Furthermore, we uncovered a sex effect, with at-risk males preferentially impaired on allocentric navigation while FH+ females were impaired on the egocentric VSTT task. This may reflect gender differences in navigational strategy, with females tending towards landmark or route navigation, and males survey-based allocentric mapping (*20*) but may also reflect sex differences in AD pathological progression, with greater parietal tau pathology in females (*19*).

Given that grid cell functioning is dependent on head direction, vestibular and optic flow information relayed to the EC via afferents from various brain regions including those affected in early AD, such as the medial temporal and parietal lobes (*25*), the PI impairment observed across hereditary and physiological risk factors for AD might reflect the unique vulnerability of the entire grid cell/PI network to disparate converging pathophysiological processes. These encompass tau deposition in the MTL, β-amyloid in the medial parietal lobe and vascular pathology, all of which are associated with family history of dementia, APOE-ε4, and CAIDE score (*15, 31, 32*). Amyloid- and tau-PET, alongside markers of vascular pathology, will help clarify the relative contributions of these differing pathologies to PI impairments.

Multimodal MRI was undertaken to identify the potential neural correlates of the PI impairments. Structural MRI using ultra high field 7T did not reveal any associations between PI and volumes of brain regions of interest, even at subfield level. This however is consistent with previous findings in presymptomatic familial AD populations (*33*) and prior PREVENT imaging studies, which did not identify clear patterns of atrophy (*34*). Considering our participant age (approximately two decades away from predicted dementia onset), the absence of volumetric change may indicate an absence of regional neurodegeneration in this cohort at this early stage in the disease process. By comparison, fMRI studies revealed an association between negative hexadirectional grid-like fMRI signal in the posterior-medial EC and PI impairment in all at-risk individuals. Further analyses aiming to understand better this negative signal in at-risk individuals revealed instead a strong unidirectional modulation of the fMRI signal. This functional imaging change may be indicative of a change in navigational strategy with increased reliance on a head direction-based approach to navigation (*35*). An overreliance on visual-based head directional signals during the outbound triangle path, at the detriment of performing accurate distance coding, could result in an angular reproduction error during the return path (*10*) – which was accentuated when distal orientation cues were removed.

In conclusion, these results indicate that impaired path integration may be the initial behavioural change in AD, prior to memory decline, and as such may represent the critical point transition from at-risk status to clinical disease onset. In addition to the benefits for clinical practice in terms of early detection and optimising future therapeutic interventions, these discoveries using a test based on the function of entorhinal cortex grid cells aids translational research in delivering a platform by which studies of AD at the cellular level may be linked to understanding the onset of the clinical disorder.

## Acknowledgements

We are extremely grateful to the participants and leadership team of the PREVENT Dementia cohort, especially Katie Wells, and staff at the Wolfson Brain Imaging Centre, MRC (Medical Research Council) Cognition and Brain Sciences Unit, Institute of Public Health and Cambridgeshire-Peterborough NHS Foundation Trust for their help in study delivery.

## Funding

Merck Investigator Studies Program grant MISP-57175 (DC)

Alzheimer’s Society grants 178, 264 and 397 (DC, CN, PREVENT Dementia)

UK National Institute for Health Research Clinical Research Network and Biomedical Research Centre Cambridge grant 1215-20014 (PREVENT Dementia, JOB), Oxford (IK), Imperial (PM)

US Alzheimer’s Association grant TriBEKa-17–519007 (PREVENT Dementia)

Alzheimer’s Research UK (DC, LS)

Wellcome grant 098436/Z/12/B (CTR), grant 202805/Z/16/Z; (NB, AC)

UK Medical Research Council grant SUAG/046 G101400 (RH), Dementia’s Platform UK grant (IK)

National Institute of Neurological Disorders and Stroke of the National Institutes of Health grant K99NS126715 (MS)

## Author contributions

Conceptualization: DC

Methodology: CN, DC, MP, RH, ZJ, MS, MED, TW, NB

Software: MS, NB, AC, TW

Investigation: CN, MP, CR, CTR

Visualization: CN, ZJ

Funding acquisition: DC, CWR

Project administration: CN, JOB, LS, IK, PM, KR, CWR

Supervision: DC, JOB, LS

Writing – original draft: CN, DC

Writing – review & editing: All

## Competing interests

The authors declare that they have no competing interests.

## Data availability

Code is available at https://tinyurl.com/yc3ybr6e and data is being made available on the open-source ADDI platform https://www.alzheimersdata.org/ (link TBC)

## Notes

### Competing Interest Statement

The authors have declared no competing interest.

