## Supplementary Materials for "Path integration selectively predicts midlife risk of Alzheimer’s disease"

Materials and Methods

S1.1 Participants

We recruited participants to this study from the PREVENT-Dementia prospective cohort (total N=701), a multi-site research programme that seeks to understand the origin point and sequence of biological and clinical changes in the pre-dementia phase of dementia diseases. The programme main entry requirements were age 40-59 and the absence of dementia or other neurological conditions at baseline, with recruitment aiming to enrich for individuals with a parental family history of dementia. The protocol has been described in detail elsewhere (*36*). Recruitment methods to this study included PREVENT newsletters and word of mouth during PREVENT Year 2 or Year 5 follow up study visits across four sites (West London, Cambridge, Oxford and Edinburgh). Power calculations derived from our study in patients with mild cognitive impairment (MCI) (*13*) were based on the primary outcome measure of our VR navigation task, Euclidean location error, for which controls had a mean (SD) of 1.27 (0.86) and patients with prodromal AD 2.55 (SD 0.92). On this basis 25 matched pairs would have allowed us to show a mean difference of 0.60 between groups; that is a mean of 1.27 in the negative group and a mean of 1.87 in the positive group with an alpha risk=0.05 and a power of 0.80.

123 potential participants initially expressed interest to take part. One potential participant was not suitable due to contraindications to using immersive virtual reality (severe motion sickness) and 22 participants later declined, resulting in 100 people giving written informed consent for their participation. Participants were stratified by APOE-ε4 status and parental family history of dementia, and did not differ on demographic variables (Table S1). When stratified by sex, given that we considered sex differences, females were moderately more educated than males (Table S7), but years of education was controlled for in all below analyses. All study team members remained blinded to participant risk statuses until recruitment and data collection was completed.

Of these 100 participants, 55 additionally gave consent to take part in a 7 Tesla MRI scan. Inclusion criteria were participation in the amyloid PET PREVENT sub-study or giving a CSF sample within the main PREVENT programme, and exclusion criteria were contraindication to scanning at 7 Tesla. This MRI subgroup was stratified on the same factors and also did not differ on demographic variables (Table S5). This study and the main PREVENT programme were performed in accordance with the Declaration of Helsinki and each was respectively approved by institutional review boards at the NHS London Camberwell St-Giles (ref. 12/LO/1023) and West London Research Ethics Committees (ref. 18/LO/2418).

S1.2 Experimental tasks

*Virtual reality path integration task*

All path integration task data was collected at the University of Cambridge, the methods of which have previously been described elsewhere (*13*). In brief, the task required participants to complete a triangle by walking between three numbered cones presented sequentially at eye-level within an open field virtual environment viewed through a head-mounted display. The open field was bordered by navigational features projected at infinity to represent boundary cues, with no local landmarks, in order to prevent use of egocentric beaconing strategies (*37*). An auditory stimulus sounded at the appearance of each cone to prompt participants towards the next cone, and cones disappeared when reached. After walking in an L-shape to reach cone three from cone one (the ‘outward path’), participants were instructed to return to their remembered location of cone one (the ‘return path’) and press a trigger on the hand-held controller. This logged their estimated location and ended the trial.

To examine the effect of supportive environmental cue availability on path integration performance, three different conditions for the return path were used. Each condition entailed a change to the environment appearance when participants reached cone three to initiate the return path: condition A, no change with all cues available; condition B, removal of surface texture; condition C, removal of distal landmarks; as described in (*13*). In addition, three different environments with varying appearance were used to maintain engagement in the task. Participants performed 12 trials per condition, totalling 36 per participant. Testing time varied from 30-60 minutes depending on participant walking speed and optional rest breaks. The order of conditions presented to participants was pseudo-randomised to remove order effects and ensure participants did not become over reliant on external allothetic vs idiothetic cues to solve the task. The locations of cones, configuration and size of triangles were also pseudo-randomised.

External base stations mapped out a 4x4m^2^ virtual test space within which participant location and task responses were tracked with a sampling rate of 0.1s to provide raw coordinate data. Triangle return path distances ranged from 3.6 to 4m to vary path integration difficulty and at least 1m of clear space boarded the test space. For safety precautions, a researcher was in close proximity at all times, and an ‘out of border’ warning message appeared in participants’ line of vision to discourage walking if they moved 30cm beyond the test space border.

Prior to the task start, participants had the opportunity to explore the environment in a short 20s habituation period and complete 5 practice trials for which feedback on performance was given. Participants were instructed to complete the task as quickly and accurately as possible, using whatever strategy they liked, but were discouraged from retracing their outward path via cone 2 to estimate where cone 1 was. No performance feedback was given during the remaining trials.

The task was administered with immersive virtual reality using the HTC Vive VR hardware system and Steam VR software. Initially this was run on the MSI VR One laptop with Intel Core i7-7820HK, 16GB RAM and GeForce GTX 1080, which was worn as a backpack to enable free, untethered participant movement during the task. However, equipment failure during data collection necessitated replacing the laptop with the Dell Desktop PC Precision 6820 Tower X-Series with Intel Core i9-10900 and GeForce RTX 2080. To enable free movement, the Vive Wireless Adaptor was additionally used.

*fMRI grid cell task*

The GC task was presented as three blocks of a 10-minute video followed by a short memory test. The video required participants to watch themselves be passively navigated through a virtual room from a first-person perspective and learn the locations of 7 target objects in the room. Target objects were every-day, household items and were highlighted by a hovering orange cone above them. Objects appeared progressively as movement within the video proceeded to cover the entire space. Movements were sequences of forward translations and rotations of varying angles. Passive participant viewing without movement control aimed to reduce motion induced through use of a joystick or button box and control the degree of room exploration per participant. The virtual room consisted of four grey walls decorated with different items to provide orientation cues, which stayed the same across all three videos – only the target objects within the room changed. When the video ended, participants were shown three images per each object from the room; one in the correct object location and two distractor images. Participants were required to select the correct image via a corresponding button on the button box, with accuracy and reaction time recorded. The next block began after the set of object location questions ended. The grid-cell-like representation (GC) task was programmed in Unity software (V2018.2.9f1) and was presented on an MRI compatible LCD screen that participants viewed through a mirror mounted on the head coil at an angle of 14°. Before scanning, participants were given verbal instructions and shown a three-minute practice video of the task. Participant position and heading direction was sampled roughly every 20ms during the video which enabled us to approximate grid event timestamps, with all participants viewing the same video per block (Section 1.5).

*MRI data acquisition*

MRI data were acquired on a 7T Terra MR system (Siemens, Erlangen, Germany) with a Nova Medical 1Tx/32Rx head coil at the Wolfson Brain Imaging Centre, University of Cambridge. Sequences were aligned to the 7TUK harmonisation protocol (*38*). First, a whole brain T1-weighted MP2RAGE volume was obtained (TR 3500ms, TE 2.58ms, TI 1 = 725ms, TI 2 = 2159ms, resolution 0.7mm isotropic, GRAPPA = 3, matrix size = 224 x 224 x 157, FA 1 = 5°, FA 2 = 2°). Second, a high-resolution partial T2-weighted structural volume was obtained with slices orientated perpendicular to hippocampal long axis (TR 8080ms, TE 76ms, resolution 0.4mm x 0.4mm, slice thickness 1mm + 10% gap, GRAPPA = 2, matrix size = 224 x 224 x 54, FA = 60°). Finally, the fMRI session was run using T2*-weighted gradient echo planar images (EPIs) with an in-plane resolution of 1.5mm x 1.5mm (42 axial slices, TR 2531ms, TE 22ms, slice thickness 1mm, GRAPPA = 2, FA = 73°, matrix size = 192 x 192 x 42). Slices were centred on and orientated parallel to the hippocampus long axis. Higher between plane resolution aimed to minimise dropout due to partial-volume effects, while lower within plane resolution aimed to increase signal-to-noise ratio. Total scan time was 75 minutes.

*MRI pre-processing*

T1-weighted structural images were produced via offline PSIR (phase-sensitive inversion recovery) reconstruction on all MP2RAGE data. T1-weighted and T2-weighted images were manually inspected for artefacts and bias field corrected using the N4 algorithm (*39*). The EPI series were preprocessed using FSL v5.0.8 (FMRIB, Oxford, UK) and SPM12 (<http://www.fil.ion.ucl.ac.uk/spm/>) in MATLAB 2019a (Mathworks, MA, USA). First, images were manually inspected for artefacts and corrected for distortions using reverse phase encoded method via FSL-topup (*40*). EPIs then underwent motion correction with SPM realign, reslice, and slice time correction to account for differences in interleaved slice acquisition times. All analyses were carried out on native space images to prevent potential signal distortions during nonlinear normalisation to a common space. All images (EPIs T1-weighted and T2-weighted) were coregistered using the default ANTs linear transformations (*41*). In cases of registration failure, images were manually coregistered in ITKSNAP (*42*).

*ROI definition*

Regions-of-interest (ROIs) were selected *a priori* either due to their hypothesised role in navigation functions or early susceptibility to pathology in initial AD stages. These included the whole and posterior-medial entorhinal cortex, hippocampal subfields (subiculum and CA1), retrosplenial cortex and posterior-cingulate cortex. For non-medial temporal lobe ROIs, T1-weighted images underwent normalisation to MNI305 atlas space, brain extraction, tissue segmentation (CSF, grey matter, white matter), and parcellation according to the Desikan-Killiany atlas using the Freesurfer image analysis suite (v7.1.0, <https://surfer.nmr.mgh.harvard.edu/>). Pial surface misplacements and erroneous WM segmentation were manually corrected on a slice-by-slice basis if individual brain processing failed. The isthmus cingulate ROI was used as a proxy retrosplenial cortex ROI mask following previous work (*13*).

Medial-temporal ROIs were created in the subject’s T2-weighted space using a semi-automatic approach with the Automatic Segmentation of Hippocampal Subfields (ASHS) software V2.0 (*43*) and IKND Magdeburg 7T multi-template atlas (*44*). ASHS outputs per hemisphere per participant were manually inspected and corrected. This included erroneously included CSF and meninges voxels or misplaced grey/white matter or anterior/posterior borders judged using established heuristic rules (*43*, *44*). Specifically for the posterior-medial entorhinal cortex, which is not available in ASHS, a publicly available common-space mask based on DTI connectivity (*45*) was used to manually trace posterior-medial entorhinal cortex voxels visible on the high-resolution T2 using ITK-SNAP (*42*). The mask was warped into individual native space via i) average group T1-weighted template made using the opensource toolkit Advanced Normalisation Tools ANTs (v2.3.4) diffeomorphic template construction algorithm (*46*) and ii) individual participant T1 and T2-weighted images, all co-registered and transformed using ANTs (*41*). In some participants the warped common space posterior-medial entorhinal mask extended anteriorly beyond the field of view of the T2-weighted image; these voxels were not included in the final ROI mask. Manual segmentation was performed by two raters with an average Dice similarity coefficient across all ROIs for 5 subjects of 0.92, indicating good inter-rater reliability.

S1.3 PREVENT Data Acquisition

Risk factor statuses and control cognitive data were collected as part the main PREVENT Dementia Program at each site (*14*, *36*).

**Family history status.** Participants self-reported parental family history status with details on dementia type and age of diagnosis if applicable and known. Reported parental dementia types from participant history were 86% either Alzheimer’s disease or Alzheimer’s disease with mixed vascular pathology, 5% Parkinson’s or Lewy Body Dementia, and the remaining 9% unknown. Participants were classified as FH+ if they had positive histories on either maternal, paternal or both sides. For estimated years to onset of dementia calculations, if both parents were diagnosed, the parent with the earlier onset age was used.

**APOE-ε4 genotyping.** Genomic DNA was isolated from whole blood samples collected during PREVENT visits and genotyping was performed using the TaqMan polymerase chain reaction (PCR)-based method, previously described in (*21*). Participants were classed as APOE-ε4 + if they carried either one or two allele copies.

**CAIDE dementia risk score.** The Cardiovascular risk factors, Ageing and Incidence of Dementia (CAIDE) DRS is a physiological-based risk scoring tool derived from a prospective cohort study that identified weighted variables in midlife predictive of future dementia (*47*). It has since been validated in additional populations against CSF and neuroimaging measures, with finalised variables including age, sex, education, hypertension, cholesterol, physical activity levels, and body mass index. In this study, CAIDE score was used without APOE-ε4 status to examine the effect of predominantly modifiable risk factors on navigation performance. It was used as a continuous variable on a scale of 0-15, but for analyses with both continuous predictor and response variables, CAIDE was considered as a binary variable of either below or above the median CAIDE score. Statistical analysis with the CAIDE did not involve controls for age, sex and education as they are included within the score. Cohort baseline measures were used to calculate scores.

**Comparator cognitive assessments.** During PREVENT visits, participants completed the digital COGNITO test battery (*22*) and additional stand-alone cognitive assessments to assess global cognitive function. In this study, measures of allocentric and egocentric processing, episodic and visual association memory were selected from these PREVENT assessments to compare against path integration, for which the administrative procedures and task details have previously been described (Table S3).

S1.4 Data Analyses

Statistics were performed in R v4.0.4 using the lme4 (*48*) and emmeans (*49*) packages. Differences between participant demographics based on risk factor and sex stratification were assessed via t-tests or non-parametric Mann-Whitney U test for continuous variables (depending on the normality of the data), and Chi-square test for categorical variables. Where appropriate in the following sections, all model residuals were inspected for deviations from homoscedasticity and normality.

*Path integration task*

All outcome measures and variables were extracted and calculated in MATLAB 2019a (The MathWorks In., Massachusetts). Each trial was manually inspected for data integrity. Trials where participants adopted a ‘retracing’ strategy or did not initiate the return path were excluded (0.6% of total trials). Trials where participants went beyond the virtual test space boundary (‘out of bounds’) were also excluded (34.6% of total trials), in line with previous work (*13*). These trials were qualitatively different to normal trials, as participants received an extra spatial cue informing their current position when the boundary was reached. We used chi-square tests to assess if proportions of out of bounds trials differed between stratified groups (see Section S2.1).

Following previous research (*50*)(*51*), the primary outcome measure for the path integration task was Location Error in virtual metres, reflecting the Euclidean distance between estimated and actual locations of cone one. We calculated distances using Eq.1, with coordinates of cone one estimated (X_1_,Y_1_) and true (X_2_,Y_2_) locations for Location Error:

Eq.1	$Distance= \sqrt{\left( X_{1}-X_{2} \right)^{2}+ \left( Y_{1}-Y_{2} \right)^{2}}$

First, an interaction effect of all risk factors, return condition type and sex on location error was assessed via a mixed linear model. We chose mixed modelling given the clustered and incomplete nature of the data (12 trials per one of three return conditions, with out of bounds trials excluded), in line with previous path integration literature (*8*, *13*). Covariates included age, years of education, and a random intercept of i) trial order number and ii) unique participant identifier with random slopes of return condition type to assess for participant variance across repeated trials. A chi-square test was used to assess for differences in the number of out of bounds trials between stratified risk groups.

In a second analysis we used multiple linear regression models to explore the change in average performance across conditions based on interactions of condition type in the mixed model results. We separately predicted change in location error between baseline and no optic flow conditions and between baseline and no distal cues conditions; in each case mean baseline performance was subtracted from the other conditions per participant to derive the outcome measure. An interaction effect of all risk factors and sex on this change in mean location error was examined. Covariates included age and years of education. Interaction effects were tested with ANOVA tests and post-hoc contrasted pairwise using t-tests Tukey-corrected for multiple comparisons.

Two additional outcome measures of Absolute Angular and Distance Error decomposed the Location Error into linear and rotational error contributions. Distance error reflects the accuracy of participant distance estimation of the return path length and was determined using the absolute values of *D_true_* – *D_estimated_*, where *D_true_* refers to the true distance between cone 3 and cone 1, and *D_estimated_* the participants’ estimated distance between cone 3 and their triggered position. Angular error reflects the accuracy of participant rotation at cone three to return to cone one and was determined using the absolute values of *A_true_* – *A_estimated_*, where *A_true_* refers to the true rotation angle from cone 3 and cone 1, and *A_estimated_* the participants’ estimated angle between cone 3 and their triggered position. Each angle ($\theta$) was calculated using Eq.2:

Eq.2	$\theta=atan2d(\vec{v1} \times\vec{v2}, \vec{v1} . \vec{v2})$

where $\vec{v1}$ represents the trajectory vector between cones two and three, $\vec{v2}$ the vector between cone three and either the true or estimated location of cone one. atan2d is a Matlab function that takes the arctangent (in degrees) of the cross and dot product of two vectors to derive the angle between them. Another series of mixed linear models with same covariates and predictors were used to assess the effect of risk factors and sex on change in absolute angular and distance error.

We created a signed allocentric angular error outcome measure to assess the directionality of angular errors. The angular difference was mapped in the interval $[-{180}^{o}, {180}^{o}]$ using the Matlab wrapto180 function as $wrapTo180(A_{true}-A_{estimated})$. Therefore, negative values indicate over-turning in the allocentric point of view, whereas positive values indicate under-turning in the allocentric point of view. For instance, if $A_{true}={120}^{o}$ and $A_{estimated}=-{220}^{o}$, then the signed allocentric angular error will be $-{20}^{o}$ (an over-turning of ${20}^{o}$), whereas the absolute angular error (described above) is ${340}^{o}$. We additionally calculated this for out of bounds trials by taking the position of the boundary collision as a proxy measure of initial angular estimate for the return path. This enabled us to circumvent the data bias of overturning errors introduced by excluding out of bounds trials (see S2.1).

Finally, we used ANOVA nested model comparison to explore the proportion of variance explained by different risk factors. We repeated the above linear regression models on change in performance from baseline to “no distal cues” using individual risk factors interacting with sex, controlled for age and education. We conducted F-tests to compare adjusted R^2^ between each of these models against the full model used above.

*Comparator cognitive assessments*

We used multiple linear regression models to explore interactive effects of all risk factors and sex on task performance for the comparator assessments, in keeping with the path integration analysis. We additionally ran separate linear regression models per individual risk factor. Covariates included age, years of education and PREVENT visit date to confirm that differences in time-locking across visit dates for PREVENT cognitive assessments and the path integration study participation did not affect results. To compare the relative predictive value of path integration to other assessments, we performed cross-validated, logistic regression using elastic-net regularisation to optimise the Area-Under-the-Curve (AUC) of the Receiver-Operator Characteristic (ROC). We predicted a “double-risk” status (FH+/APOE-ε4 + vs any other combination, based on earlier model performance) using performance on the VR path integration task (viz. change in location error from baseline to “no distal cues” condition) plus performance on the other tasks explored above, as well as age, sex, and education. We used 1000 random permutations of training vs test.

*MRI volumes*

For the MTL ROIs, regional T2-weighted volumes were extracted using the Insight Toolkit Convert3D software ([www.itksnap.org/c3d](http://www.itksnap.org/c3d)). T1-weighted isthmus cingulate and posterior cingulate regional volumes, as well as T1-weighted total intracranial volume, were extracted using Freesurfer v7.1.0 (as described above). To reduce multiple comparisons, volumes from the left and right hemisphere were summed, and to correct for variations in brain size, volumes were expressed as a percentage of total intracranial volume to estimate relative grey matter. We ran an additional analysis using raw volumes with total intracranial volume as a covariate to confirm this analysis approach did not alter results.

First, we used multiple linear regressions predicting each individual ROI per each individual risk factor (FH, APOE-ε4, CAIDE) to assess effects of risk on brain structure. We then used a three-way interaction between each ROI volume, risk factor status and sex to predict change in location error across conditions to assess effects of risk on brain-behaviour relationships. All models had covariates of age and years in education. The false discovery rate method was used to correct for planned multiple ROI comparisons per each risk factor and response variable (*52*).

Finally, to establish if multivariate contributions of all ROIs better explained structural brain-behaviour relationships, we used a multiple regression with all ROIs as predictors with age, sex, and education covariates.

*fMRI grid-cell-like representations*

Putative measures of grid cell codes were extracted from the pre-processed EPIs using a reproducible, standardised approach with the GridCAT v1.04 (*27*) and CircStat (*53*) toolboxes in MATLAB (2017b, The MathWorks In., Massachusetts) and SPM12 (<http://www.fil.ion.ucl.ac.uk/spm/>). The spatial properties of the grid patterns and the non-linearity of the mapping from location through neural activity to the macroscopic blood-oxygen-level-dependent (BOLD) fMRI signal suggest that grid population activity could be detectable in fMRI. Namely, the orientations of the grid-firing patterns of both local and distant grid cells are clustered, despite differences in grid scale (*54*, *55*). Thus, the different neural firing dynamics when running in directions aligned versus misaligned to grid axes (i.e. some cells firing a lot, others a little, versus all cells firing an intermediate amount) would generate different fMRI signal strengths (*56*).

Data analysis here used two general linear models with parametric modulators of *sin* and *cos* to i) estimate individuals’ grid orientation (in 60° space) using the time-varying translation events in half of the data, and ii) derive measures of model fit using this estimated orientation for modelling translation events in the remaining data. Data were partitioned using odd/even translation events and nuisance regressors for head movement were included in the general linear models. Only voxels masked by the right posterior-medial entorhinal subdivision mask were used to calculate outcome metrics, in line with previous findings (*7*, *8*, *56*). The primary outcome measure was grid-cell-like representation magnitude, where higher magnitude entailed better fit of the general linear model for the estimated mean grid angle (*27*). Secondary measures of between-voxel orientation coherence and within-voxel orientation coherence over time respectively provided measures of spatial and temporal stability. Spatial stability was calculated using Rayleigh’s test for non-uniformity of circular data using voxelwise mean grid orientations. Temporal stability was calculated by comparing orientation values within voxels between the first and second half of each scanning run, expressed as the percentage of voxels with a change of less than 15° in orientation. We included these secondary measures because temporal but not spatial stability was demonstrated to be the cause of low magnitude grid codes in young APOE-ε4 carriers relative to non-carriers (*7*).

The fMRI grid outcome metrics were normally distributed, so one-sample t-tests were used to assess if magnitudes and spatial stabilities of grid cell like representations were significantly different from zero and if temporal stability was significantly different from chance (50%) across all participants. Differences in metrics between FH/APOE-ε4 risk groups and sex were compared using Welch’s two-sample t-tests given unequal variances while associations between metrics and age or CAIDE lifestyle risk score were calculated with Pearson correlations.

We used multiple linear regression models to test if i) metrics predicted change in location error from baseline to no distal cues conditions and following on from this ii) if a three-way interaction between grid-cell-like representation magnitude, risk status and sex modulated this relationship. We used covariates of age and education, and sex in the first regression models. To assess the continuous x continuous CAIDE interaction with grid-activity magnitude, we used Johnson-Neyman intervals to explore the range of CAIDE values for which the association between grid activity and path integration performance was significant.

We additionally performed a range of control analyses (see Supplementary Text). Grid cell like representation magnitudes were calculated using either 4-fold, 5-fold or 7-fold rotational symmetry (instead of expected 6-fold) to confirm the specificity of the grid-characteristic patterns. Temporal signal-to-noise ratio for the posterior-medial entorhinal cortex was calculated by dividing voxel-wise mean timeseries by its standard deviation, and this was associated against grid-cell-like representation magnitudes using Pearson correlation.

Supplementary Text

S2.1 Out of bounds trial exclusion

We labelled trials as ‘out of bounds’ if participants tracked 30cm beyond the 4x4m^2^ test area which resulted in a safety warning message appearing in their sightline telling them to stop walking. As the message provided a local egocentric landmark cue that could have disrupted path integration-based computations, these trials were excluded from analysis of location error. While APOE-ε4 + showed more frequent out of bounds events of borderline significance (χ^2^(1,3564) = 4.00, *p* = 0.05), FH+ or individuals with above median CAIDE score did not (both *p* > 0.16), suggesting minimal bias was introduced by excluding out of bounds trials.

We explored whether excluding out of bounds trials affected our main conclusions about overturning errors by developing a proxy measure of angular error for out of bounds trials. We used the location of the boundary collision instead of location of participant trigger pull to provide an estimate of participants’ initial heading direction for out of bounds trials. Because boundary collisions were mostly caused by under-turning (Fig. S2A), we used signed angular error in allocentric space to better account for the form of error made. This confirmed that overturning was still the predominant form of angular error event with inclusion of out of bounds trials (Fig. S2B).

We also used this all-trial signed angular error to assess if inclusion of out of bounds trials influenced our main conclusions about the effect of midlife AD risk and sex on path integration performance, which was based on regular trials only. Re-running the same analysis on all trials (namely, a multiple linear regression predicting an interaction between all risk statuses and sex on the change in signed angular error from baseline to no distal cues) showed an overall similar but weaker pattern of results (CAIDE *F*_1,78_ = 4.03, *p* = 0.048; FH x APOE-Ε4 *F*_1,78_ = 2.20, *p* = 0.142; FH x sex *F*_1,78_ = 3.86, *p* = 0.053; Fig. S3). However, because the location of boundary collision did not reflect the true final participants’ estimate of cone 1, we did not use this proxy outcome measure to draw conclusions about the data.

S2.2 Grid-like activity controls

Hexadirectional grid-cell-like representation activity magnitude predicted path integration performance (as described in main text), but population-level grid-like representation activity magnitude across all participants was non-significant, i.e. not greater than zero (one-tail t-test *t*_52_=0.5, *p*=0.6). However, stratifying participants by above and below median path integration performance groups showed there was significantly higher grid cell representation magnitudes in the better performing group (two sample t-test *t*_47_ = 2.00, *p* = 0.04) and the better performing group trended towards significant grid activity representations (mean = 0.109; one-tail t-test *t*_26_ = 0.8, *p* = 0.200).

Temporal stability was not significantly different from chance (one-way t-test *t*_52_ = 0.04, *p* = 0.500) but there was significant spatial stability (mean Rayleigh Z = 2.38, one-way t-test *t*_52_=12.0, *p*<0.001). There was no effect of FH+ or APOE-ε4+ on temporal stability (Welch two-way t-tests, both *p* > 0.10), but there was a trend association between increased CAIDE and decreased temporal stability (Pearson’s *r* = -0.24, *p* = 0.090), in line with findings in APOE-ε4 carriers and older adults (*7*, *26*). Grid-cell-like activity magnitude was not related to the volume (*r*=0.012, *p*=0.930) or temporal signal-to-noise ratio (*r*=-0.042, *p*=0.760) of the posterior-medial entorhinal cortex. There was no effect of risk status on temporal signal-to-noise ratio (Welch two-sample t-test, all *p* > 0.2) or volume of the posterior-medial entorhinal cortex (all *p* > 0.3).

To confirm the specificity of the hexadirectional grid cell-like activity, we repeated the same multiple linear regressions predicting change in location error using standard controls of 4, 5 and 7-fold symmetrical models of grid cell activity. This revealed no significant associations (4-fold β = 0.04 ± 0.10, t_46_ = 0.45, *p* = 0.650; 5-fold β < 0.01 ± 0.08, t_46_ < 0.01, *p* = 0.999; 7-fold β = 0.12 ± 0.11, t_46_ = 1.12, *p* = 0.270).

S2.3 Negative hexadirectional grid-like activity

In an exploratory analysis we examined for unidirectional modulation of the pmEC grid signal, hypothesising that it might underlie the negative hexadirectional grid signal observed in at-risk individuals with poorer PI performance. If a strong unidirectional signal is present it could interfere with the estimation of the grid angle, thereby resulting in a signal that is significant but rotated between the estimation and test data sets, hence a negative overall magnitude. Such a unidirectional signal might reflect increased activity of head-direction like processing, which aligns with the impaired angular estimation during PI without visual orientation cues in at-risk individuals. Namely, if they were over-relying on head direction signals from visual cues in the environment during the outbound path of the triangle, and therefore did not perform accurate integration of distance, it could result in angular error during the return path (*10*). To this end, we found that stronger unidirectional grid-like fMRI signals predicted an increased decline in PI with removal of distal visual orientation cues (β = 0.31 ± 0.09, *t_46_* = 3.50, *p* = 0.001; Fig. 3E). To further explore a head-direction like effect on the signal, we examined if there was a clustering of estimated mean grid orientations of the unidirectional modulated signal in individuals using the Rayleigh test for uniformity of directions. While this was not significant for the population overall, it was significant in males only (Z = 3.34, *p* = 0.030; females Z=0.745, *p* = 0.479), in line with the apparent stronger association of unidirectional signal and PI impairments observed in males (Fig. 3F). The average mean orientation was 181º (Fig. S4), consistent with the use of a single salient landmark visible on a room wall during the fMRI task. Future work is needed to confirm the association with AD risk factors.

| Table S1. Demographics for all participants stratified either by family history or APOE-Ε4 status | | | | | | | |
| --- | --- | --- | --- | --- | --- | --- | --- |
| Characteristic | Family history positive  N = 62 | Family history negative  N = 38 | *p* | APOE-ε4 positive  N = 32 | APOE-ε4 negative  N = 66 | *p* | Whole sample  N=100 |
| Sex |  |  |  |  |  |  |  |
| Female (%) | 41 (66%) | 23 (61%) | *0.70^a^* | 19 (59%) | 43 (65%) | *0.70^a^* | 64 (64%) |
| Age |  |  |  |  |  |  |  |
| Mean yrs | 57.1 ± 5.12 | 55.8 ± 5.45 | *0.20* | 55.2 ± 5.46 | 57.3 ± 5.10 | *0.07* | 56.6 ± 5.25 |
| Education |  |  |  |  |  |  |  |
| Mean yrs | 17.0 ± 3.10 | 16.8 ± 2.86 | *0.70* | 16.8 ± 2.86 | 16.9 ± 2.91 | *0.60^b^* | 16.9 ± 3.00 |
| APOE-ε4  Positive (%)  NA (%) | 18 (29%)  2 (3%) | 14 (37%)  0 | *0.60^a^* | -  - | -  - |  | 32 (32%)  2 (2%) |
| Family history ^†^   Positive (%) | - | - |  | 18 (56%) | 42 (64%) | *0.60^a^* | 62 (62%) |
| Family history type  Maternal (%)  Paternal (%)  Both (%) | 28 (45%)  43 (69%)  9 (15%) | -  -  - |  | 10 (56%)  11 (61%)  3 (17%) | 16 (38%)  32 (76%)  6 (14%) | *0.50^a^*  *0.30^a^*  *1.00^a^* | -  -  - |
| CAIDE  Mean score   NA (%) | 5.14 ± 2.19  3 (3%) | 4.87 ± 2.23  0 | *0.30^b^* | 4.94 ± 2.33  0 | 5.08 ± 2.15  1 (2%) | *0.50^b^* | 5.03 ± 2.20  3 (3%) |
| a Pearson Chi-square test; b Wilcoxon rank sum test; ^†^ parental; CAIDE = Cardiovascular risk factors, Ageing and Dementia Incidence Study. | | | | | | | |

| Table S2. ANOVA nested model comparison of predictors for change in location error from baseline to no distal cues conditions | |
| --- | --- |
| Predictors | **Adjusted R^2^** |
| FH x APOE x CAIDE x sex | 0.26 |
| FH x APOE x sex | 0.20 |
| FH x APOE x CAIDE | 0.13 |
| FH x sex | 0.11 |
| APOE x sex | 0.07 |
| CAIDE x sex | 0.05 |
| All models controlled for age and education, and sex were not already included. FH = family history; APOE = APOE-ε4 allele, CAIDE = Cardiovascular Risk Factors, Aging and Dementia Study dementia risk score | |

| Table S3. Comparator cognitive assessments used in PREVENT. | | |
| --- | --- | --- |
| Exam | **Metrics used** | **Relevant cognitive domain** |
| Addenbrookes Cognitive Exam III (ACE) | Total score /100 | Screening for global impairment |
| Visual short-term binding test (VSTBT) (*6*) | A’ of shape-colour binding performance | Non-verbal frontal and medial temporal associative memory |
| Four Mountains Task (4MT)  (*24*) | Total score /15 | Hippocampal allocentric spatial memory |
| Virtual Reality Supermarket Trolley Task (VRSTT) (*23*) | Total score /20 | Retrosplenial egocentric spatial memory |
| COGNITO Name-Face Association (NameFace) (*22*) | Total score /9 | Non-verbal medial-temporal paired associative memory |
| COGNITO Narrative recall (NarRecall) (*22*) | Total score /27 | Verbal medial-temporal episodic memory |

| Table S4. Effect of AD risk factors on comparator task performance | | | | | | |
| --- | --- | --- | --- | --- | --- | --- |
| Risk Factor | **4MT** | **VRSTT** | **VSTBT** | **Narrative Recall** | **Name-Face** | **ACE-R** |
| FH+ | 10.1 ± 0.3 | 9.8 ± 0.2 | 0.73 ± 0.03 | 13.6 ± 0.6 | 5.4 ± 0.2 | 96.7 ± 0.4 |
| FH- | 11.4 ± 0.4 | 10.1 ± 0.3 | 0.71 ± 0.03 | 12.6 ± 0.7 | 5.5 ± 0.3 | 96.4 ± 0.5 |
| *p _FH_* | **0.016** | ns | ns | ns | ns | ns |
| *p _FH x sex_* | ns | **0.008 ^B^** | ns | ns | ns | ns |
| APOE-ε4 + | 10.6 ± 0.3 | 10.3 ± 0.3 | 0.69 ± 0.03 | 13.3 ± 0.8 | 5.15 ± 0.3 | 96.9 ± 0.5 |
| APOE-ε4 - | 10.7 ± 0.5 | 9.6 ± 0.2 | 0.74 ± 0.02 | 13.2 ± 0.6 | 5.55 ± 0.2 | 96.5 ± 0.4 |
| *p _APOE_* | ns | ns | ns | ns | ns | ns |
| *p _APOE-Ε4 x sex_* | ns | ns | ns | ns | ns | ns |
| CAIDE β | -0.21 ± 0.18 | 0.07 ^C^ | -0.01 ± 0.01 | 0.11 ± 0.31 | -0.39 ± 0.09 | -0.07 ^C^ |
| *p* | ns | ns | ns | ns | **< 0.001** | ns |
| FH+/APOE-ε4 + | 9.7 ± 0.6 | 10.2 ± 0.4 | 0.69 ± 0.04 | 14.2 ± 1.0 | 5.27 ± 0.4 | 97.3 ± 0.4 |
| FH-/APOE-ε4 + | 12.0 ± 0.6 | 10.5 ± 0.5 | 0.70 ± 0.05 | 12.4 ± 1.2 | 5.01 ± 0.5 | 96.3 ± 0.8 |
| FH+/APOE-ε4 - | 10.1 ± 0.5 | 9.3 ± 0.4 | 0.75 ± 0.03 | 13.3 ± 0.7 | 5.34 ± 0.3 | 96.2 ± 0.5 |
| FH-/APOE-ε4 - | 11.2 ± 0.5 | 10.1 ± 0.4 | 0.71 ± 0.04 | 12.9 ± 0.9 | 5.84 ± 0.4 | 96.5 ± 0.6 |
| *p _Multifactor status_* | **0.019 ^A^** | ns | ns | ns | ns | ns |
| *p* values shown corrected for age, sex, and education in multiple linear regression per individual risk factor. A = driven by singular difference in FH+/APOE-ε4 + vs FH-/APOE-ε4 + (*p_Tukey_* = 0.058). B = driven by difference in FH+ vs FH- in females (*p* = 0.006). C = Spearman’s Rho shown given ceiling effects in the data. FH = family history; CAIDE = Cardiovascular risk factors, Ageing and Dementia Incidence Study Dementia Risk Score. | | | | | | |

| Table S5. Demographics for MRI subgroup, stratified either by family history or APOE-ε4 status | | | | | | | |
| --- | --- | --- | --- | --- | --- | --- | --- |
| Characteristic | Family history positive  N = 34 | Family history negative  N = 21 | *p* | APOE-ε4 positive  N = 18 | APOE-ε4 negative  N = 36 | *p* | Whole sample  N=55 |
| Sex |  |  |  |  |  |  |  |
| Female (%) | 18 (53%) | 12 (57%) | *0.70^a^* | 11 (61%) | 18 (50%) | *0.60^a^* | 30 (55%) |
| Age |  |  |  |  |  |  |  |
| Mean yrs (SD) | 57.2 ± 5.02 | 56.2 ± 4.33 | *0.37* | 55.9 ± 4.71 | 57.2 ± 4.80 | *0.30^b^* | 56.9 ± 4.75 |
| Education |  |  |  |  |  |  |  |
| Mean yrs (SD) | 16.6 ± 3.05 | 16.5 ± 3.06 | *0.92* | 16.7 ± 3.06 | 16.6 ± 2.94 | *1^b^* | 16.5 ± 3.02 |
| APOE-ε4   Positive (%)   NA (%) | 12 (35%)  1 (3%) | 6 (28%)  0 | *0.80^a^* | -  - | -  - |  | 18 (33%)  1 (2%) |
| Family history ^†^   Positive (%) | - | - |  | 12 (67%) | 21 (58%) | *0.80^a^* | 34 (62%) |
| Family history type   Maternal (%)   Paternal (%)   Both (%) | 14 (41%)  23 (67%)  3 (9%) | -  -  - |  | 7 (39%)  7 (39%)  2 (11%) | 6(17%)  16 (44%)  1 (3%) | *0.10^a^*  *0.90^a^*  *0.50^a^* | -  -  - |
| CAIDE   Mean score (SD)   NA (%) | 5.72 ± 2.17  2 (6%) | 5.05 ± 2.31  0 | *0.20^b^* | 5.61 ± 2.34  0 | 5.37 ± 2.34  1 (3%) | *0.80^b^* | 5.45 ± 2.23  3 (3%) |
| a Pearson Chi-square test; b Wilcoxon rank sum test; ^†^ parental; CAIDE = Cardiovascular risk factors, Ageing and Dementia Incidence Study. | | | | | | | |

| Table S6. AD risk factor differences on relative ROI volumes (% of total intracranial volume) | | | | | | | | | | | |
| --- | --- | --- | --- | --- | --- | --- | --- | --- | --- | --- | --- |
| Risk Factor | **CA1** | **CA2** | **CA3** | **DG** | **Sub** | **pmEC** | **alEC** | **PrC** | **PhC** | **RSC** | **PCC** |
| FH+ | 0.142 | 0.006 | 0.029 | 0.101 | 0.206 | 0.089 | 0.094 | 0.106 | 0.092 | 0.476 | 0.567 |
| FH- | 0.146 | 0.007 | 0.029 | 0.100 | 0.206 | 0.082 | 0.096 | 0.103 | 0.100 | 0.479 | 0.584 |
| *p* | 0.35 | 0.14 | 0.84 | 0.68 | 0.72 | 0.38 | 0.57 | 0.78 | 0.11 | 0.88 | 0.32 |
| APOE-ε4 + | 0.138 | 0.006 | 0.028 | 0.095 | 0.204 | 0.089 | 0.092 | 0.107 | 0.094 | 0.478 | 0.564 |
| APOE-ε4 - | 0.147 | 0.007 | 0.029 | 0.103 | 0.207 | 0.084 | 0.097 | 0.104 | 0.096 | 0.477 | 0.579 |
| *p* | 0.11 | **0.04** | 0.50 | **0.02** | 0.75 | 0.47 | 0.66 | 0.33 | 0.58 | 0.88 | 0.59 |
| CAIDE β | < -.001 | < -.001 | < -.001 | < -.001 | 0.003 | < -.001 | < -.001 | < -.001 | < .001 | .007 | < .001 |
| *p* | 0.68 | **0.05** | 0.16 | 0.88 | **0.02** | 0.69 | 0.93 | 0.31 | 0.64 | **0.08** | 0.96 |
| *p* values shown corrected for age, sex, and education in multiple linear regression but not multiple comparisons; all *p_FDR_* > 0.183. β regression values for CAIDE predictor. FH = family history; CAIDE = Cardiovascular risk factors, Ageing and Dementia Incidence Study Dementia Risk Score; CA = cornu ammonis, DG = detate gyrus, Sub = subiculum, pmEC = posterior-medial entorhinal cortex, alEC = anterior-lateral entorhinal cortex, PrC = perirhinal cortex, PhC = parahippocamapal cortex, RSC = retrosplenial cortex, PCC = postering cingulate cortex. | | | | | | | | | | | |

| Table S7. Demographics for main sample, stratified by sex | | | |
| --- | --- | --- | --- |
| Characteristic | Male  N = 36 | Female  N = 64 | *p* |
| Age |  |  |  |
| Mean yrs | 57.6 ± 5.16 | 56.0 ± 5.27 | *0.20* |
| Education |  |  |  |
| Mean yrs | 16.1 ± 2.39 | 17.4 ± 3.22 | *0.04** |
| APOE-ε4   Positive (%)   NA (%) | 13 (36%)  0 | 19 (29%)  2 (3%) | *0.70^a^* |
| Family history ^†^   Positive (%) | 21 (58%) | 41 (64%) | *0.70^a^* |
| a Pearson Chi-square test; b Wilcoxon rank sum test; ^†^ parental; CAIDE = Cardiovascular risk factors, Ageing and Dementia Incidence Study. | | | |

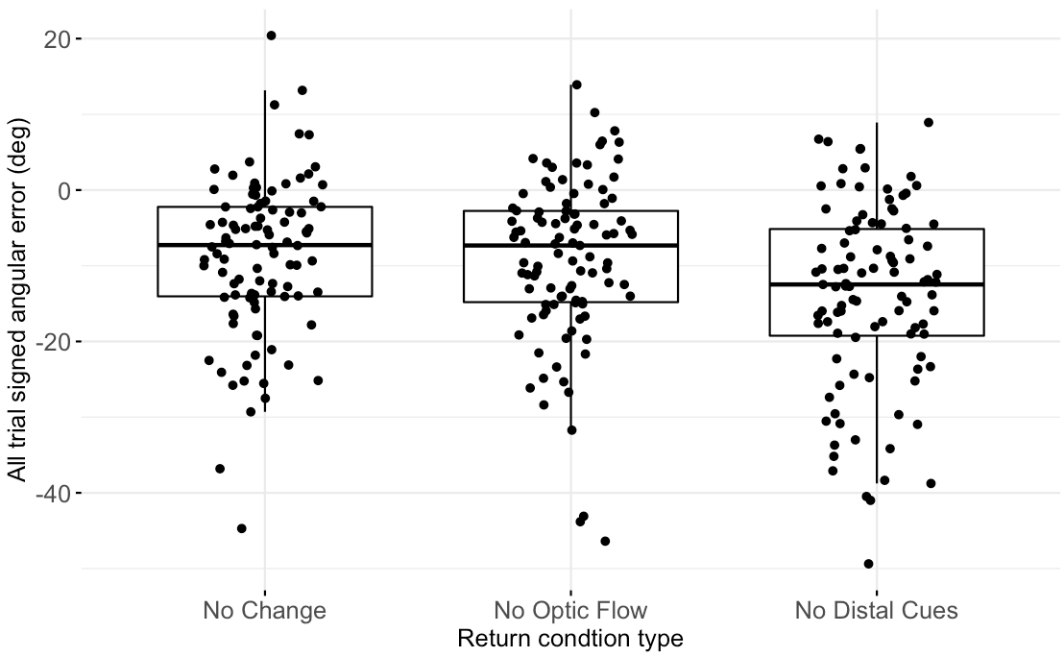
**Figure S1.** Change in allocentric signed angular error from baseline to “no distal cues” conditions

All participants showed negative angular errors, corresponding to over-turning, which was magnified when distal cues were removed.

**Figure S2.** Allocentric signed angular error for regular trials and out of bounds trials.

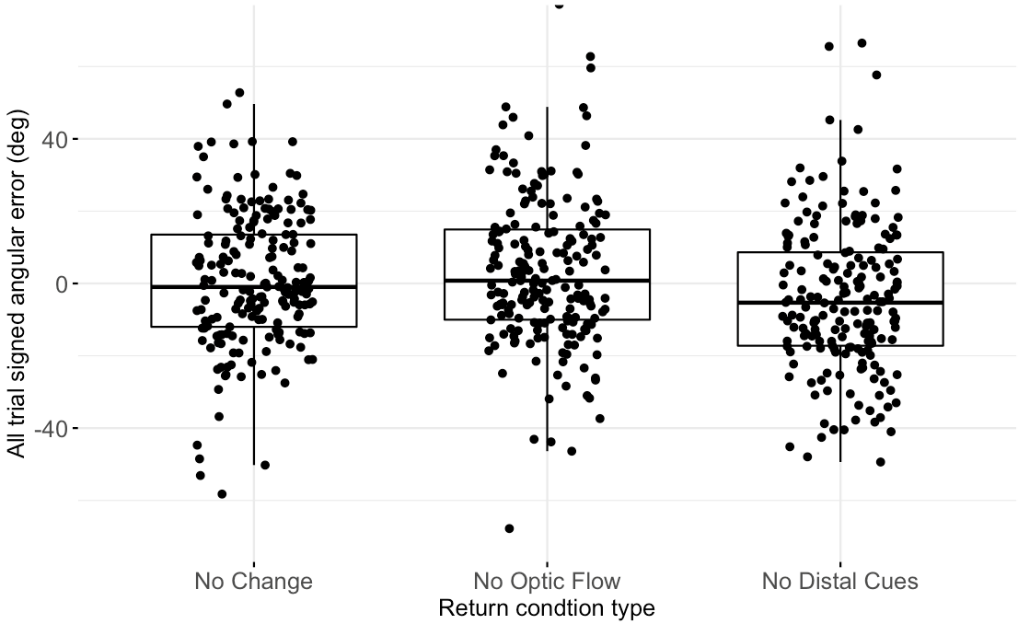

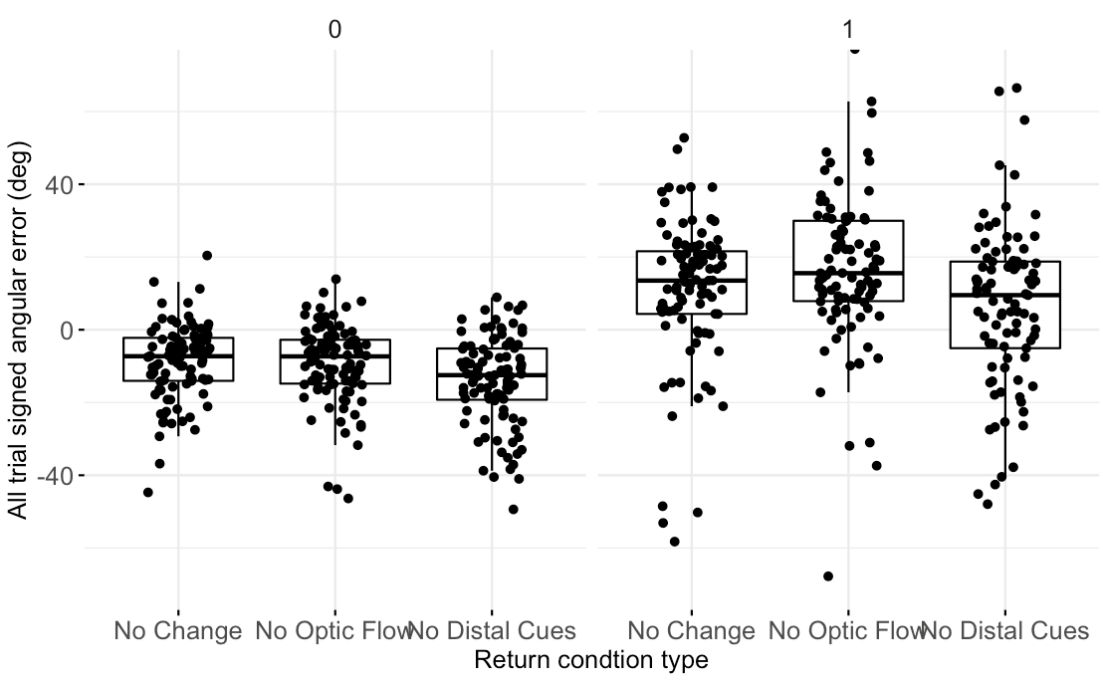

A

B

Regular trials

Out of bounds trials

(**A**) Out of bounds trials associated with more under-turning (positive angular errors) than over-turning (negative values). (**B**) Overall across all trials, both regular and out of bounds, the angular errors (especially for the no distal cues trials) were still driven by over-turning even with out of bounds trials accounted for.

**Figure S3.** Replicating main effects after inclusion of out of bounds trials with proxy angular error measure.

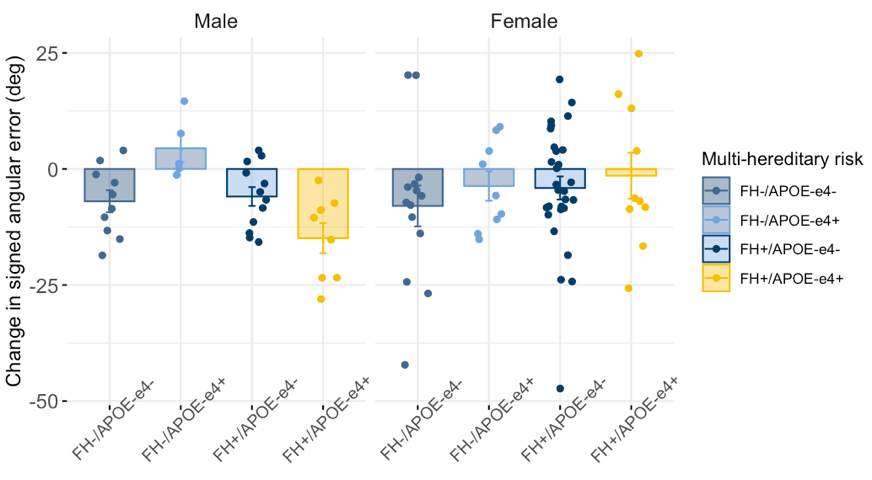

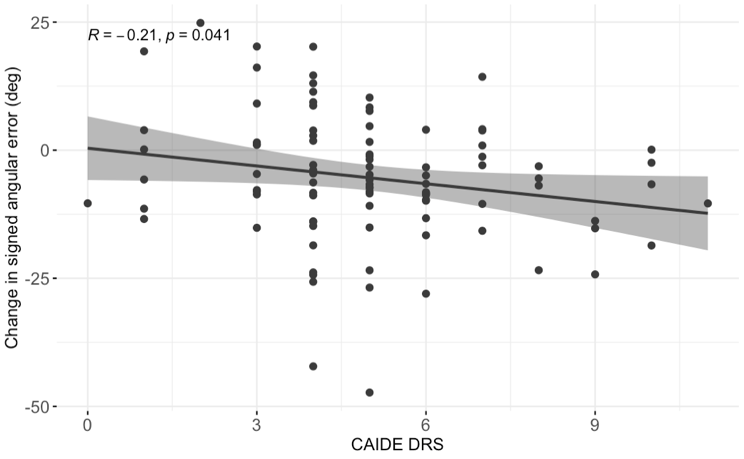

A

B

(**A**): male FH+/APOE-ε4+ visually trend towards larger over-turns across all trials from baseline to no distal cues condition, recapitulating the main effects seen for regular trials only. (**B**): higher CAIDE associates with greater over-turning errors from baseline to no distal cues conditions. Negative values indicate increases in over-turning relative to cone 1; positive values increases in under-turning.

**Figure S4.** Mean estimated grid orientations of the unidirectional signal are significantly clustered in males only.

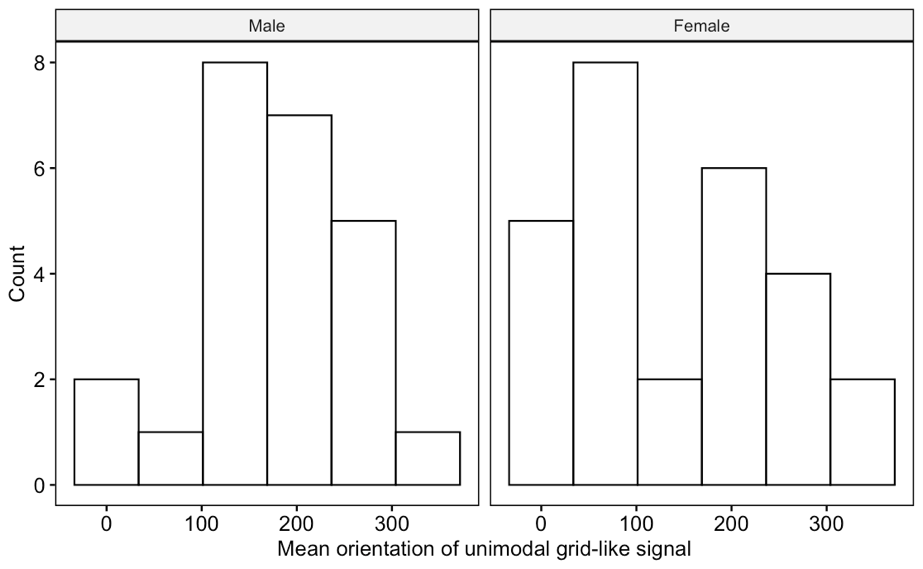

Left: male individuals show a clustering of estimated mean orientations around 181º, whereas females show uniformly distributed orientations (right).
